# A Fixed Moderate-dose Combination of Tiletamine+Zolazepam Outperforms Midazolam in Induction of Short-term Immobilization of Ball Pythons (*Python regius*)

**DOI:** 10.1101/341438

**Authors:** Lynn J. Miller, David P. Fetterer, Nicole L. Lackemeyer, Matthew G. Lackemeyer, Ginger C. Donnelly, Jesse T. Steffens, Sean A. Van Tongeren, Jimmy O. Fiallos, Joshua L. Moore, Shannon T. Marko, Luis A. Lugo-Roman, Greg Fedewa, Joseph L. DeRisi, Jens H. Kuhn, Scott J. Stahl

**Affiliations:** United States Army Medical Research Institute of Infectious Diseases, Fort Detrick, MD, USA; Integrated Research Facility at Fort Detrick, National Institute of Allergy and Infectious Diseases, National Institutes of Health, Fort Detrick, Frederick, MD, USA; Department of Biochemistry and Biophysics, University of California San Francisco, San Francisco, CA, USA; Stahl Exotic Animal Veterinary Services, Fairfax, VA, USA

**Author notes:** Correspondence and requests for materials should be addressed to • L.J.M., Veterinary Medicine Division, United States Army Medical Research Institute of Infectious Diseases (USAMRIID), 1425 Porter Street, Fort Detrick, Frederick, MD 21702, USA; -xxx; -xxx, • J.H.K., Integrated Research Facility at Fort Detrick (IRF-Frederick), Division of Clinical Research (DCR), National Institute of Allergy and Infectious Diseases (NIAID), National Institutes of Health (NIH), B-8200 Research Plaza, Fort Detrick, Frederick, MD 21702, USA.

## Abstract

Laboratory animals are commonly anesthetized to prevent pain and distress and to provide safe handling. Anesthesia procedures are well-developed for common laboratory mammals, but not as well established in reptiles. We assessed the performance of intramuscularly injected tiletamine (dissociative anesthetic) and zolazepam (benzodiazepine sedative) in fixed combination (2 mg/kg and 3 mg/kg) in comparison to 2 mg/kg of midazolam (benzodiazepine sedative) in ball pythons (*Python regius*). We measured heart and respiratory rates and quantified induction parameters (i.e., time to loss of righting reflex, time to loss of withdrawal reflex) and recovery parameters (i.e., time to regain righting reflex, withdrawal reflex, normal behavior). Mild decreases in heart and respiratory rates (median decrease of <10 beats per minute and <5 breaths per minute) were observed for most time points among all three anesthetic dose groups. No statistically significant difference between the median time to loss of righting reflex was observed among animals of any group (*p* = 0.783). However, the withdrawal reflex was lost in all snakes receiving 3mg/kg of tiletamine+zolazepam but not in all animals of the other two groups (*p* = 0.0004). In addition, the time for animals to regain the righting reflex and resume normal behavior was longer in the drug combination dose groups compared to the midazolam group (*p* = 0.0055). Our results indicate that midazolam is an adequate sedative for ball pythons but does not suffice to achieve reliable immobilization or anesthesia, whereas tiletamine+zolazepam achieves short-term anesthesia in a dose-dependent manner.

## INTRODUCTION

A good understanding of the natural history, evolutionary development, and reproduction of snakes (Reptilia: Squamata: Serpentes) is paramount for many reasons including conservation efforts [1-4], evaluation of snake-specific sensory anatomy for chemoreception and behavior [5-8], and to help develop medical countermeasures against venomous snake bites, and finally the development of candidate therapeutics for human health issues including clotting disorders and cancer [9-12]. Snakes are also increasingly investigated as natural or accidental hosts for a variety of viruses [13-23], and there is increased interest in snake fungal diseases such as those caused by *Ophidiomyces ophiodiicola* [24] and the *Chrysosporium* anamorph of *Nannizziopsis vriesii* complex (CANV) [25]. Snake researchers face unique challenges for husbandry, animal restraint, biosample collection, and surgery. Varying levels of immobilization can be required depending on the procedure to be performed, the size and temperament of the snakes, or their ability to produce venom [26-28]. These challenges can be exacerbated when snakes are suspected to be, or are infected with potentially zoonotic microorganisms or viruses, and when the safety of the animal handler, and prevention of bites, becomes a primary concern.

Ball pythons (Boidae: *Python regius* Shaw, 1802) are common pets in the US but are still rarely used in experimental research [29, 30]. These snakes have gained attention among virologists as they harbor unique viruses [31-34]. Ball python nidovirus (*Nidovirales*: *Toroviriniae*) was recently identified as a likely cause for respiratory disease in this species [14, 35, 36]. Additionally, Golden Gate virus, a reptarenavirus (*Arenaviridae*: *Reptarenavirus*) was recently discovered in boa constrictors (Boidae: *Boa constrictor* Linnaeus, 1758) and annulated trees boas (Boidae: *Corallus annulatus* Cope, 1875). Golden Gate virus is the likely cause of inclusion body disease (IBD) of captive boid snakes [37-39] and is highly virulent for ball pythons [16].

Due to the increased interest in virus research involving ball pythons, we designed this study to identify a reliable method of immobilization in these snakes that permits experimental infections and sample acquisition in the absence of animal suffering. We therefore aimed to identify an anesthetic that can be injected intramuscularly without intubation (straightforward and relatively uncomplicated procedure). Administration of this anesthetic ought to lead to reliable immobilization of all study animals for at least 10–15 min with minimal depression of the cardiovascular and respiratory systems and a recovery period of less than 3 hours. Based on previously published literature for other reptiles [40-44], we hypothesized that a low, 2 mg/kg, dose of tiletamine (dissociative anesthetic) plus zolazepam (benzodiazepine sedative) would best meet these anesthesia criteria in infectious disease laboratory settings (biosafety levels 2 and higher). Here, we describe an *in vivo* comparison of low, 2-mg/kg, tiletamine+zolazepam intramuscular dosing to mid-range, 3-mg/kg, dosing of the same compound combination and high, 2-mg/kg, dosing of midazolam, a commonly used but also understudied sedative for reptiles [45-52]. Our results indicate that intramuscular administration of 3 mg/kg of tiletamine+zolazepam to ball pythons results in anesthesia that lasts sufficiently long for animal exposure to infectious agents and acquisition of biosamples.

## MATERIALS AND METHODS

### Ethics Statement

Research was conducted under an Institutional Animal Care and Use Committee (IACUC)-approved protocol at the United States Army Medical Research Institute of Infectious Diseases (USAMRIID) in Frederick, Maryland, USA. USAMRIID is accredited by the AAALAC International and adheres to principles stated in the Guide for the Care and Use of Laboratory Animals, National Research Council (2011).

### Animals

Six adult (4 female and 2 male) ball pythons (*Python regius* Shaw, 1802) were obtained from a commercial breeder, Outback Reptiles (Manassas, VA, USA), and four (3 female and 1 male) adult ball pythons were obtained from two private collectors coordinated by Stahl Exotic Animal Veterinary Services (Fairfax, VA, USA). The mean body weight of these animals was 1.57 kg ± 0.43 (range 0.80–2.29 kg). Animals were maintained in a commercially available rack system (Freedom Breeder, Turlock, CA; Model: Reptile 1010) designed to house snakes of this weight. Snakes were maintained within an appropriate optimum temperature zone. In the cage, ambient daytime temperatures were maintained at 25.5–27.5 °C, ambient nighttime temperatures were maintained at 24.0–26.7 °C, and basking spots were maintained at 31.0–35.0 °C. Heat was provided by a custom-made under-tank heating element controlled via a thermostat (Freedom Breeder). The animal room itself was maintained at 23.9 ± 1°C. Humidity was maintained at 45–70 %, and a 12-hour light/dark cycle was maintained in the room housing the racks system. All snakes were acclimatized for 7 days after arrival at the facility and were physically examined and evaluated via baseline complete blood count (CBC) and chemistry panel to ascertain health. Ball python nidovirus and reptarenavirus infection of the animals was excluded via next-generation sequencing analysis of total RNA extracted from blood samples taken from each snake as previously described [14]. Prior to the study, each snake was offered a freshly euthanized laboratory mouse (strain CD-1, Envigo, Fredrick, MD, USA) of appropriate size every 10–14 days. To avoid complications with regurgitation and improper digestion, snakes were held off feed for at least 5 days before and after anesthesia procedures were conducted.

### Baseline Ethogram

To create a baseline ethogram for each animal, normal behavior, reactiveness, and temperament were recorded for 10 minutes twice daily (between 6 and 8 am EST, and between 4 and 6 pm EST) for 7 days. A designated set of stimuli was always provided at each observation: cages were slowly opened exposing 50–70% of inner cage surface, cage furniture was calmly manipulated to conduct daily husbandry activities (e.g., changing water, checking bedding substrate for urates or feces and removal thereof), and snakes were physically stimulated by steady digital pressure against the lateral surface of the body. A summary of the behaviors noted for 1 week is documented in Table 1. This baseline ethogram was used for comparison of snake behavior pre- and post-drug administration since variation in typical reactivity levels was observed between individual snakes depending on the stimulus provided.

**Table 1.**
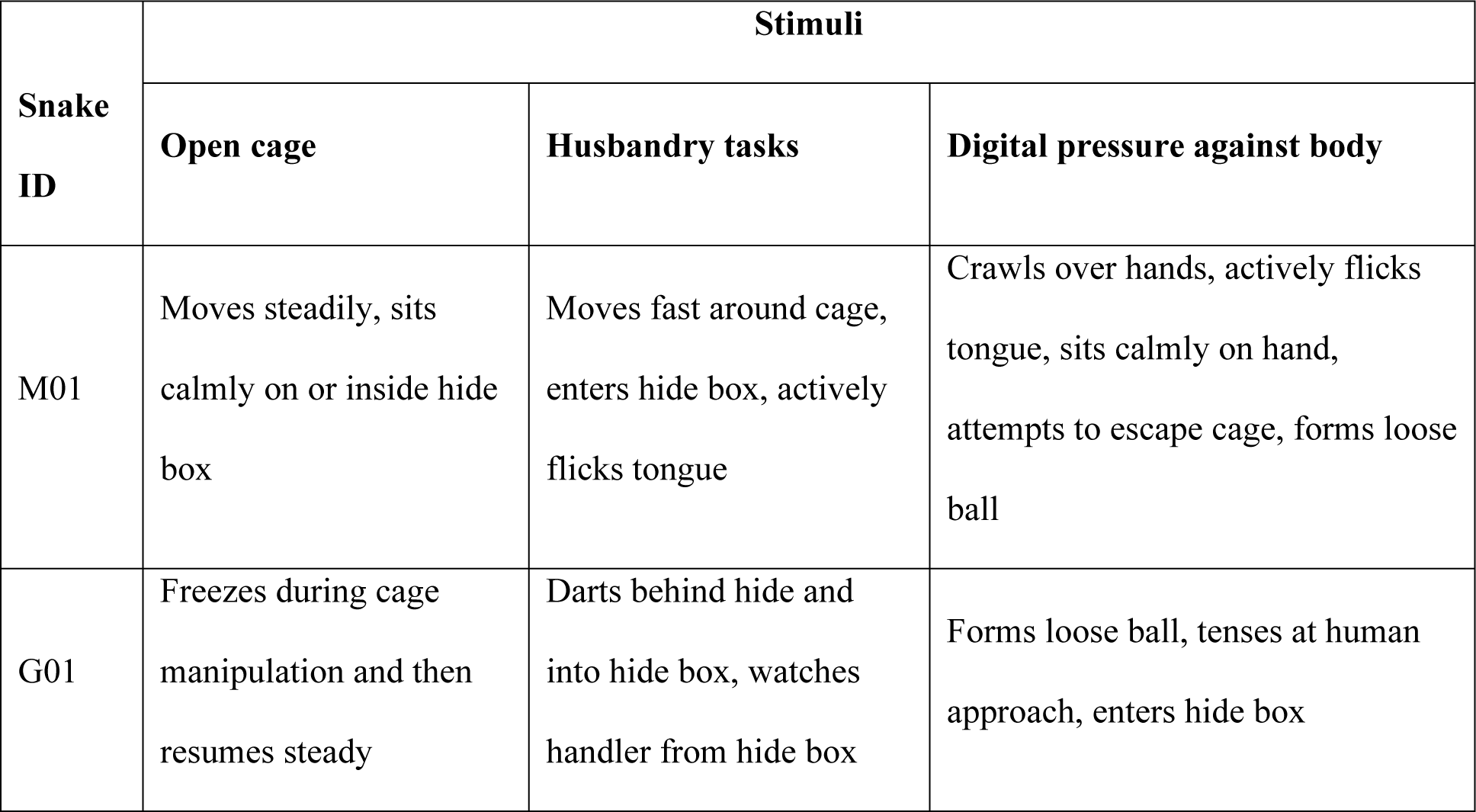
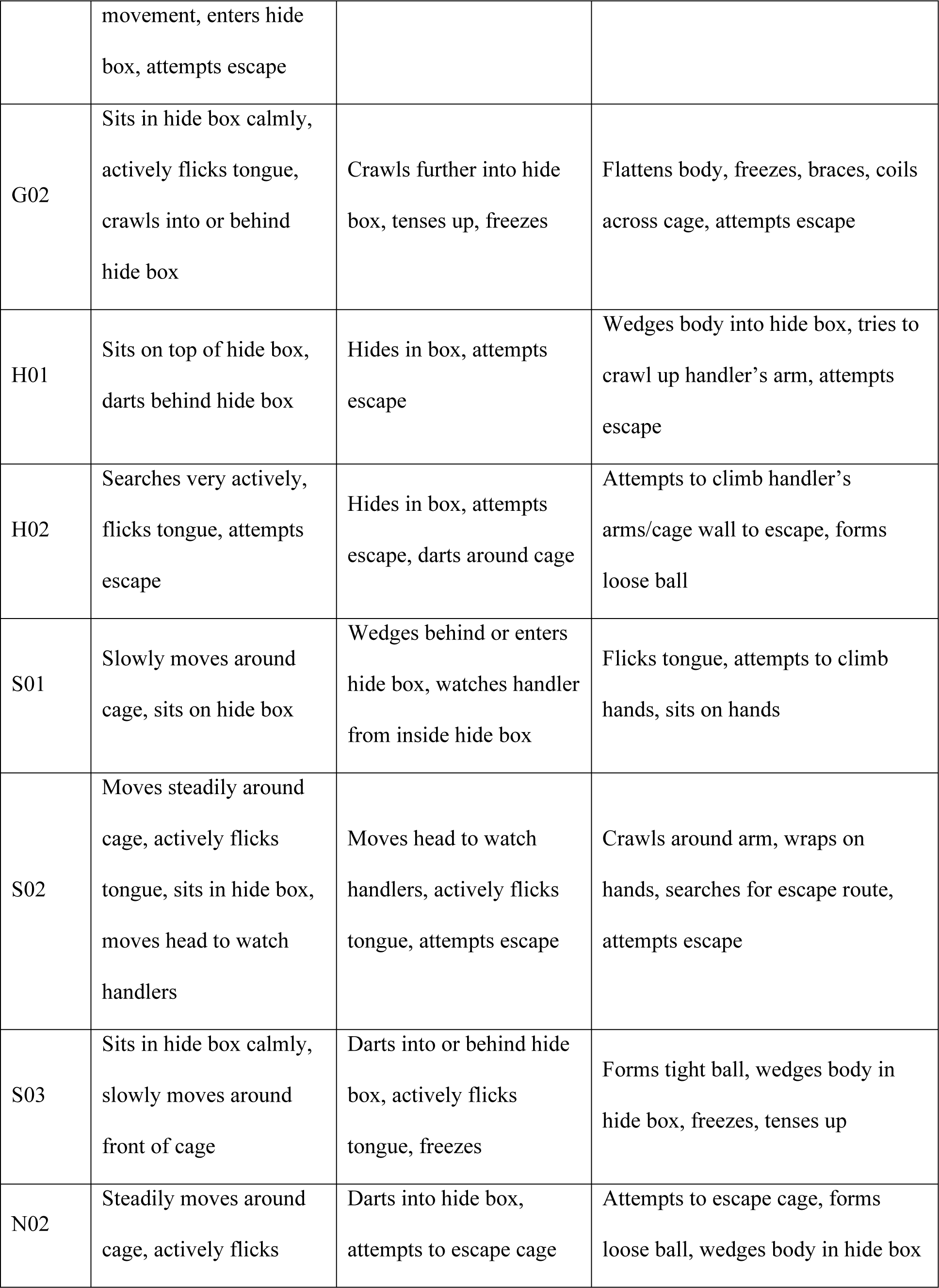
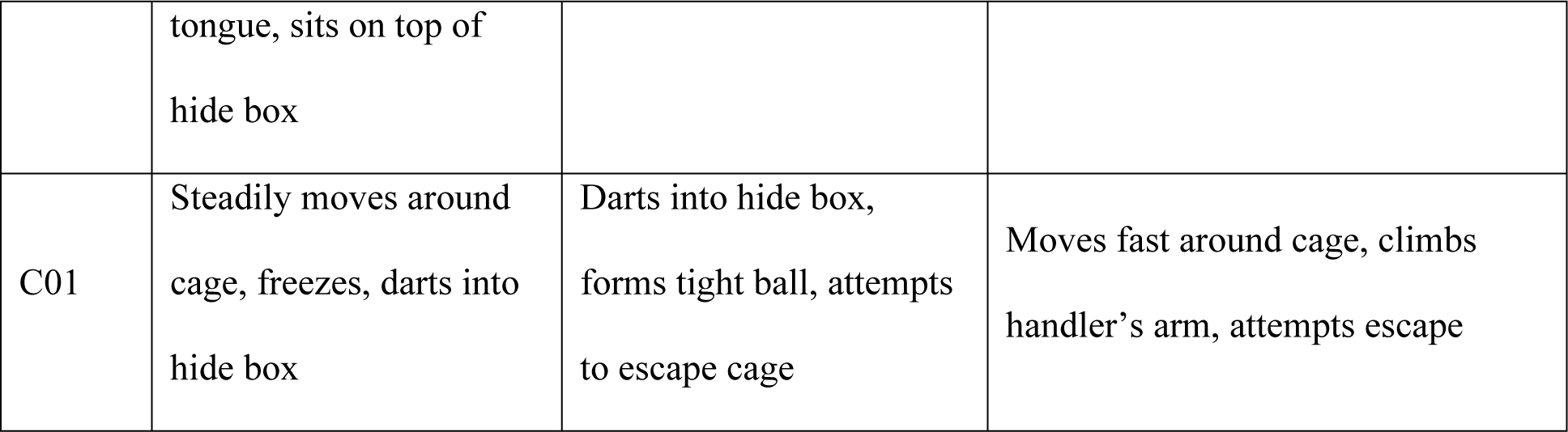
Ball Python Baseline Ethogram.

| Snake ID | Stimuli |  |  |
| --- | --- | --- | --- |
|  | Open cage | Husbandry tasks | Digital pressure against body |
| M01 | Moves steadily, sits calmly on or inside hide box | Moves fast around cage, enters hide box, actively flicks tongue | Crawls over hands, actively flicks tongue, sits calmly on hand, attempts to escape cage, forms loose ball |
| G01 | Freezes during cage manipulation and then resumes steady movement, enters hide box, attempts escape | Darts behind hide and into hide box, watches handler from hide box | Forms loose ball, tenses at human approach, enters hide box |
| G02 | Sits in hide box calmly, actively flicks tongue, crawls into or behind hide box | Crawls further into hide box, tenses up, freezes | Flattens body, freezes, braces, coils across cage, attempts escape |
| H01 | Sits on top of hide box, darts behind hide box | Hides in box, attempts escape | Wedges body into hide box, tries to crawl up handler's arm, attempts escape |
| H02 | Searches very actively, flicks tongue, attempts escape | Hides in box, attempts escape, darts around cage | Attempts to climb handler's arms/cage wall to escape, forms loose ball |
| S01 | Slowly moves around cage, sits on hide box | Wedges behind or enters hide box, watches handler from inside hide box | Flicks tongue, attempts to climb hands, sits on hands |
| S02 | Moves steadily around cage, actively flicks tongue, sits in hide box, moves head to watch handlers | Moves head to watch handlers, actively flicks tongue, attempts escape | Crawls around arm, wraps on hands, searches for escape route, attempts escape |
| S03 | Sits in hide box calmly, slowly moves around front of cage | Darts into or behind hide box, actively flicks tongue, freezes | Forms tight ball, wedges body in hide box, freezes, tenses up |
| N02 | Steadily moves around cage, actively flicks tongue, sits on top of<br>hide box | Darts into hide box, attempts to escape cage | Attempts to escape cage, forms loose ball, wedges body in hide box |
| C01 | Steadily moves around<br>cage, freezes, darts into<br>hide box | Darts into hide box,<br>forms tight ball, attempts<br>to escape cage | Moves fast around cage, climbs<br>handler's arm, attempts escape |

### Anesthesia

Snakes were assigned to three anesthesia groups in a crossover study design. The treatment sequence and order of drug administration each snake received were randomized using PROC PLAN (SAS Institute, Cary, North Carolina, USA; Version 9.4). Seven days elapsed between anesthesia trials to allow clearance of drugs. Midazolam, 5 mg/mL, was obtained from Hospira Pharmaceuticals (Lake Forest, IL, USA). Tiletamine+zolazepam, 10 mg/mL, was obtained from Zoetis LLC (Parsippany, NJ, USA). The three drug regimens tested were midazolam (2 mg/kg), tiletamine+zolazepam (2 mg/kg), and tiletamine+zolazepam (3 mg/kg). Drugs were administered to one snake at a time under restraint by two animal handlers. Administration occurred intramuscularly into paravertebral muscles located in the cranial third of each snake’s body using tuberculin needles (BD Medical, Franklin Lakes, NJ, USA), 25-gauge, 1.59 cm, 1 ml). Snakes were returned to the rack system immediately after injection for observation.

### Monitoring

Anesthetists were blinded to the drug each snake received to reduce observer bias, and multiple anesthetists covered the monitoring required for simultaneous dosing of all ten animals. Anesthetic depth was measured by evaluating the presence or absence of the righting reflex (turning the snakes on their backs and observing whether they assume normal body position thereafter) and withdrawal reflexes (observing whether the snakes attempt to move their tails after the anesthetist firmly pressured the tails for at least 3 seconds using his/her finger tips). Heart rates were measured using non-directional Doppler ultrasound system and a 9-mHz flat probe (Parks Medical Electronics, INC, Aloha, Oregon, USA). Respiratory rates were measured by visual examination of spontaneous respiration. Both physiological parameters were measured every 5–7 min after drug administration until the snakes regained righting reflexes.

During induction and recovery, animals were maintained on the basking side of their enclosures for thermal support. Entry into Stage 1 of anesthesia was defined as “exposed to the drug.” Induction time was defined as the time from administration of the drug to the time an evaluated snake lost its righting reflex. The time from the loss of righting reflex to regaining the reflex was designated as Stage 2 of anesthesia. During Stage 2 of anesthesia, the snake was immobile and minimally resistant to handling. The time passing from drug administration to loss of the withdrawal reflex was recorded as well. The anesthetic period, Stage 3 of anesthesia, was defined as the time from the loss of the withdrawal reflex to regaining the reflex. This stage was used to acquire blood and other biosamples.

Resumption of typical behavior was defined as the time from initial drug dosing to exhibiting fully alert behaviors in line with the baseline ethogram (e.g., actively investigating cage environment, investigating animal handlers, and reacting to manipulation of cage furniture). After animals regained righting reflexes, behaviors were assessed every 15–20 min.

### Statistical Analysis

The association between drug regimen and time to loss of righting and/or withdrawal reflexes was analyzed by a log-rank test stratified by animal and was summarized by median time among all snakes. Animals not experiencing an event were considered as right censored at the end of the observation period. These cases prevented the calculation of the upper confidence limit for the 2-mg/kg midazolam and 2-mg/kg tiletamine-zolazepam groups, and the calculation of the median for the 2-mg/kg tiletamine-zolazepam group. These values are denoted with “NC” (not calculated) in Table 2. The time to regain the righting and/or withdrawal reflex and/or normal behavior was considered only among those snakes experiencing the corresponding losses and were compared by a stratified log-rank test. The proportion of snakes ever reaching the loss of withdrawal or righting reflexes and the frequency distribution of snake anesthesia stages by time post-injection was determined using Friedman’s chi-square test. Minimum, maximum, and median heart and respiratory rates, obtained for each snake over the first 1–2 hours of study, were compared across treatment groups by one-way repeated measures ANOVA. Adjustments for multiple comparisons were not applied.

**Table 2.**
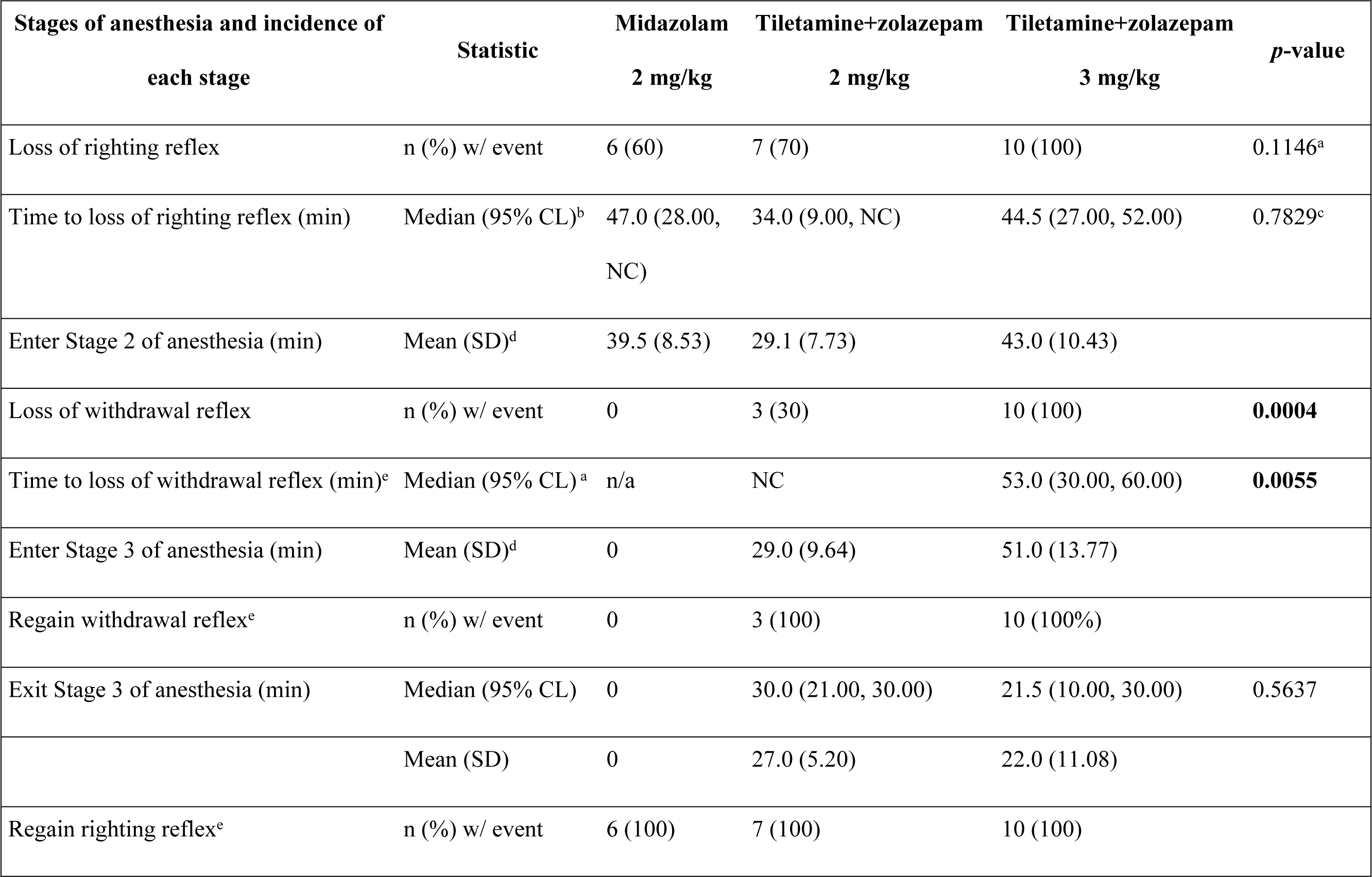
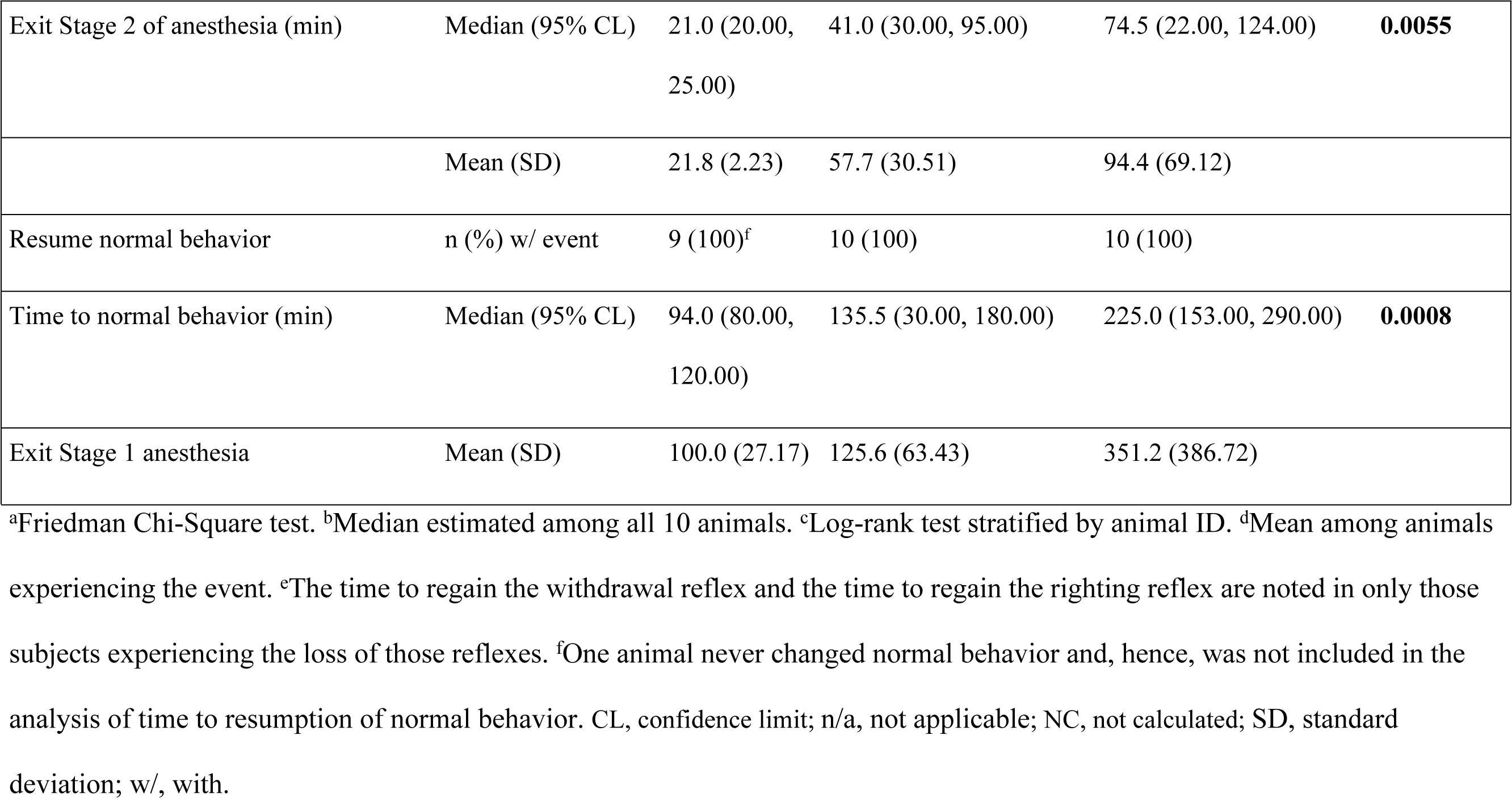
Time Course of Anesthesia Stages After Administration of Three Different Drug Regimens in Ball Pythons.

| Stages of anesthesia and incidence of<br>each stage | Statistic | Midazolam<br>2 mg/kg | Tiletamine+zolazepam<br>2 mg/kg | Tiletamine+zolazepam<br>3 mg/kg | *p*-value |
| --- | --- | --- | --- | --- | --- |
| Loss of righting reflex | n (%) w/ event | 6 (60) | 7 (70) | 10 (100) | 0.1146<sup>a</sup> |
| Time to loss of righting reflex (min) | Median (95% CL)<sup>b</sup> | 47.0 (28.00, NC) | 34.0 (9.00, NC) | 44.5 (27.00, 52.00) | 0.7829<sup>c</sup> |
| Enter Stage 2 of anesthesia (min) | Mean (SD)<sup>d</sup> | 39.5 (8.53) | 29.1 (7.73) | 43.0 (10.43) |  |
| Loss of withdrawal reflex | n (%) w/ event | 0 | 3 (30) | 10 (100) | **0.0004** |
| Time to loss of withdrawal reflex (min)<sup>e</sup> | Median (95% CL)<sup>a</sup> | n/a | NC | 53.0 (30.00, 60.00) | **0.0055** |
| Enter Stage 3 of anesthesia (min) | Mean (SD)<sup>d</sup> | 0 | 29.0 (9.64) | 51.0 (13.77) |  |
| Regain withdrawal reflex<sup>e</sup> | n (%) w/ event | 0 | 3 (100) | 10 (100%) |  |
| Exit Stage 3 of anesthesia (min) | Median (95% CL) | 0 | 30.0 (21.00, 30.00) | 21.5 (10.00, 30.00) | 0.5637 |
|  | Mean (SD) | 0 | 27.0 (5.20) | 22.0 (11.08) |  |
| Regain righting reflex<sup>e</sup> | n (%) w/ event | 6 (100) | 7 (100) | 10 (100) |  |
| Exit Stage 2 of anesthesia (min) | Median (95% CL) | 21.0 (20.00, 25.00) | 41.0 (30.00, 95.00) | 74.5 (22.00, 124.00) | **0.0055** |
|  | Mean (SD) | 21.8 (2.23) | 57.7 (30.51) | 94.4 (69.12) |  |
| Resume normal behavior | n (%) w/ event | 9 (100)<sup>f</sup> | 10 (100) | 10 (100) |  |
| Time to normal behavior (min) | Median (95% CL) | 94.0 (80.00, 120.00) | 135.5 (30.00, 180.00) | 225.0 (153.00, 290.00) | **0.0008** |
| Exit Stage 1 anesthesia | Mean (SD) | 100.0 (27.17) | 125.6 (63.43) | 351.2 (386.72) |  |

<sup>a</sup>Friedman Chi-Square test. <sup>b</sup>Median estimated among all 10 animals. <sup>c</sup>Log-rank test stratified by animal ID. <sup>d</sup>Mean among animals experiencing the event. <sup>e</sup>The time to regain the withdrawal reflex and the time to regain the righting reflex are noted in only those subjects experiencing the loss of those reflexes. <sup>f</sup>One animal never changed normal behavior and, hence, was not included in the analysis of time to resumption of normal behavior. CL, confidence limit; n/a, not applicable; NC, not calculated; SD, standard deviation; w/, with.

## RESULTS

All 10 ball pythons received one dose of each drug regimen at 7-day intervals. Administration of midazolam at 2 mg/kg or tiletamine+zolazepam at 2 mg/kg resulted in in loss of the righting reflex (Stage 2 of anesthesia, defined in Materials & Methods) in 6/10 and 7/10 snakes respectively, whereas all 10 snakes lost the righting reflex after exposure to tiletamine+zolazepam at 3 mg/kg (Table 2).

The loss of righting reflex and entry into Stage 2 of anesthesia resulted in mostly immobile and minimally resistant snakes that could be handed easily. Statistically significant differences between each anesthetic group were observed for the time each snake spent in Stage 2 of anesthesia (*p* = 0.0055): 3 mg/kg tiletamine+zolazepam resulted in a median of 74.5 min of Stage 2 anesthesia, whereas 2 mg/kg tiletamine+zolazepam and midazolam resulted in shorter times (41.0 and 21.0 min, respectively; Table 2). Statistically significant differences (*p* = 0.0004) were also observed at Stage 3 of anesthesia (defined in Materials & Methods) between the three drug regimens: only 3 mg/kg tiletamine+zolazepam resulted in the loss of the withdrawal reflex in all 10 animals (Table 2). The time to regain the withdrawal reflex was similar in all snakes that experienced loss of the withdrawal reflex independent of drug regimen. A chronological representation of the snakes’ anesthetic events and recoveries is shown in Fig. 1.

**Fig. 1.**
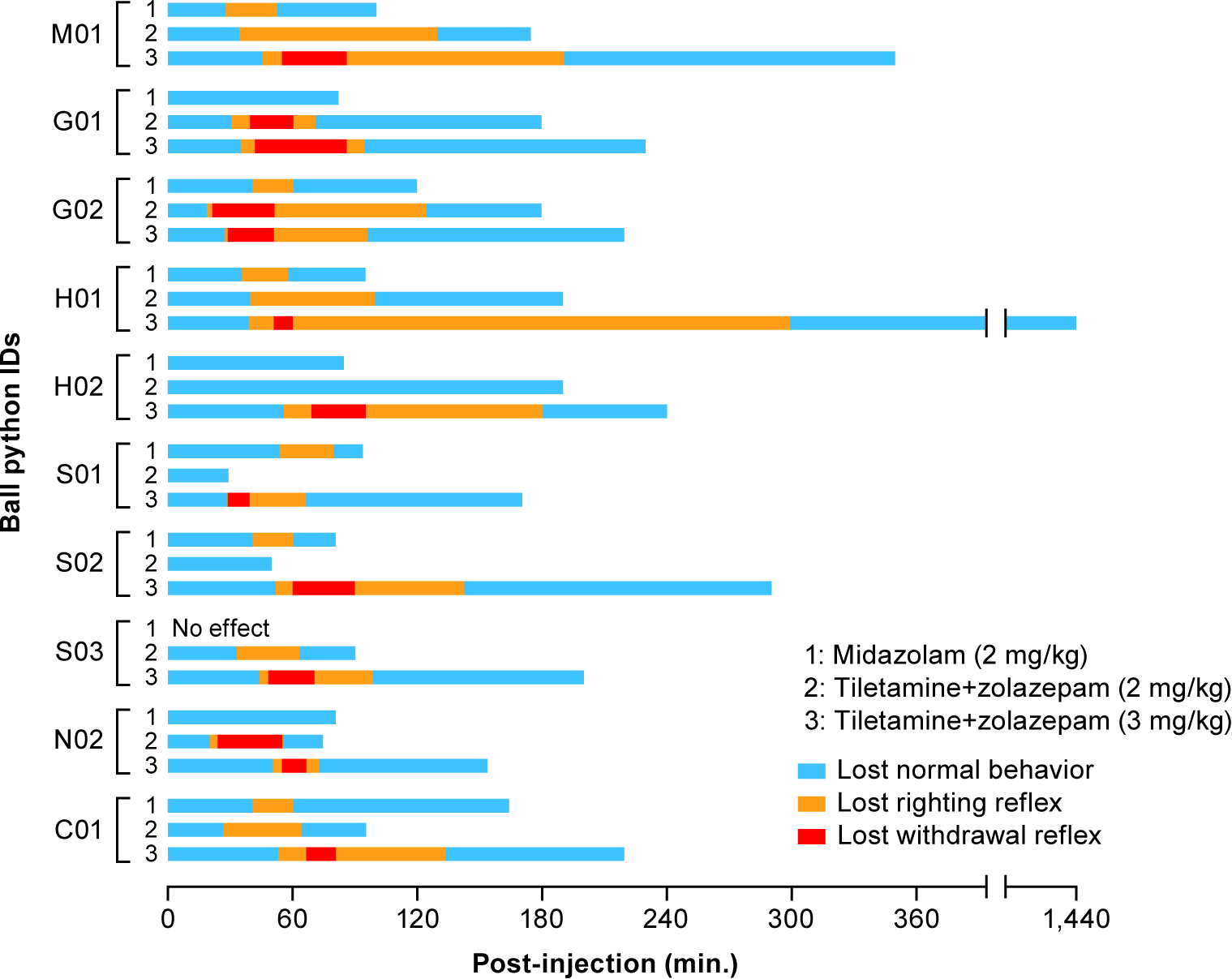
Evaluation of the Depth of Anesthesia in Ball Pythons Following Injection of Midazolam or a Fixed Combination of Tiletamine+Zolazepam. Each snake was graphed chronologically through the stages of anesthesia experienced by the animal after intramuscular injection of midazolam (2 mg/kg), tiletamine+zolazepam (2 mg/kg), or tiletamine+zolazepam (3 mg/kg). Time spent in each anesthetic stage is displayed by colored bars. The duration of anesthesia of snake H01 following administration of 3 mg/kg of tiletamine+zolazepam is denoted by an interrupted X axis. Snake H01 did not return to normal behavior until 1,440 min post-injection.

Notably, during the first 45 min after drug administration, no statistically significant differences were noted between the three drug regimens regarding the number of snakes in Stage 1 and 2 of anesthesia (Fig. 1, Table 2). Subsequently, animals exposed to 3 mg/kg of tiletamine+zolazepam took longer to regain their righting reflexes and exit Stage 2 of anesthesia compared to the 2 mg/kg group (Fig. 1, Table 2). Snakes exposed to 3 mg/kg of tiletamine+zolazepam took longer to enter Stage 3 of anesthesia (mean 51 min) than snakes exposed to 2 mg/kg of tiletamine+zolazepam (mean 29 min) and not all animals exposed to the 2 mg/kg dose lost the withdrawal reflex. However, the mean and median time needed to regain the withdrawal reflex and exit Stage 3 of anesthesia was similar between the two tiletamine+zolazepam groups (Table 2). The baseline ethogram constructed prior to study start (Table 1) was used as a definition of expected normal behaviors for each snake. A statistically significant difference was observed between the three drug regimen groups in the time to resume normal behavior. Snakes in the midazolam-treated group resumed normal behavior the fastest, followed by the 2-mg/kg and then the 3-mg/kg tiletamine+zolazepam groups (*p* = 0.0008, Table 2). No abnormal clinical signs were noted post-recovery.

Heart rate and respiratory rates were measured for 5 min for each snake to establish baselines prior to drug administration. During the first hour after drug administration, 9/10 animals experienced a significant decrease in the mean heart rate from -3.7 to -14.0 beats per minute (bpm) in the midazolam-treated group, 8/10 animals experienced a mean change of +0.2 to -15.2 bpm in the 2-mg/kg tiletamine+zolazepam-treated group, and 10/10 animals experienced a mean change of -5.55 to -13.05 bpm in the 3-mg/kg tiletamine+zolazepam-treated group (Table 3).

**Table 3.** Impact of Midazolam or Tiletamine+Zolazepam on Mean Heart and Respiratory Rates in Ball Pythons.

| Physiologic parameter | Hour(s) post-injection | Statistic | Midazolam |  |  | Tiletamine+zolazepam<br>2 mg/kg |  |  | Tiletamine+zolazepam<br>3 mg/kg |  |  | *p*-value<sup>a</sup> |
| --- | --- | --- | --- | --- | --- | --- | --- | --- | --- | --- | --- | --- |
|  |  |  | Mean | 95% CL |  | Mean | 95% CL |  | Mean | 95% CL |  |  |
| Heart rate (bpm) |  | Baseline | 68.70 | 60.57 | 76.83 | 67.30 | 60.22 | 74.38 | 67.75 | 62.95 | 72.55 | 0.9390 |
|  | 1 | Maximum | -3.7000 | -8.2625 | 0.8625 | 0.2000 | -5.8775 | 6.2775 | -5.5500 | -10.5074 | -0.5926 | 0.1260 |
|  |  | Median | -9.1000 | -14.4766 | -3.7234 | -8.1500 | -17.9223 | 1.6223 | -9.6500 | -15.7737 | -3.5263 | 0.9220 |
|  |  | Minimum | -14.0000 | -21.0897 | -6.9103 | -15.2000 | -25.2874 | -5.1126 | -13.0500 | -19.1164 | -6.9836 | 0.8062 |
|  | 2 | Maximum | 1.4000 | -3.0508 | 5.8508 | -3.1000 | -9.8986 | 3.6986 | -2.7500 | -7.9175 | 2.4175 | 0.2876 |
|  |  | Median | -2.4500 | -8.1366 | 3.2366 | -8.2000 | -17.5715 | 1.1715 | -6.8500 | -11.7195 | -1.9805 | 0.3349 |
|  |  | Minimum | -6.3000 | -12.8307 | 0.2307 | -10.8000 | -20.7455 | -0.8545 | -10.7500 | -15.7440 | -5.7560 | 0.4377 |
| Respiratory rate<br>(breaths/min) |  | Baseline | 26.15 | 20.77 | 31.53 | 21.10 | 15.48 | 26.72 | 23.70 | 20.55 | 26.85 | 0.3835 |
|  | 1 | Maximum | -0.8500 | -4.7828 | 3.0828 | 1.5000 | -0.4839 | 3.4839 | 2.4000 | -0.03642 | 4.8364 | 0.2943 |
|  |  | Median | -4.1500 | -7.6547 | -0.6453 | -1.9000 | -4.1949 | 0.3949 | -0.1500 | -2.6963 | 2.3963 | 0.1316 |

Performance of Injectable Anesthetic Agents in Ball Pythons
| Physiologic parameter | Hour(s) post-injection | Statistic | Midazolam |  |  | Tiletamine+zolazepam<br>2 mg/kg |  |  | Tiletamine+zolazepam<br>3 mg/kg |  |  | *p*-value<sup>a</sup> |
| --- | --- | --- | --- | --- | --- | --- | --- | --- | --- | --- | --- | --- |
|  |  |  | Mean | 95% CL |  | Mean | 95% CL |  | Mean | 95% CL |  |  |
|  |  | Minimum | -6.6500 | -9.9715 | -3.3285 | -5.7000 | -8.7520 | -2.6480 | -2.3000<sup>b,c</sup> | -5.0406 | 0.4406 | **0.0212** |
|  | 2 | Maximum | 0.8500 | -3.2039 | 4.9039 | 1.2000 | -3.6887 | 6.0887 | -0.5000 | -5.3547 | 4.3547 | 0.6171 |
|  |  | Median | -0.8500 | -4.0449 | 2.3449 | -1.0000 | -5.9993 | 3.9993 | -3.7000 | -8.7617 | 1.3617 | 0.3107 |
|  |  | Minimum | -2.6500 | -5.3162 | 0.01624 | -3.8000 | -9.8762 | 2.2762 | -6.0000 | -11.2784 | -0.7216 | 0.3787 |

<sup>a</sup>One-way repeated measures analysis testing the hypothesis that all three treatments are equal. <sup>b</sup>Significantly different from the
midazolam group. <sup>c</sup>Significantly different from the 2-mg/kg tiletamine+zolazepam group. bpm, beats per minute.

No significant differences were noted between groups regarding decreases in heart rates. A slight increase in heart rate was seen in 2/10 animals of the 2-mg/kg tiletamine+zolazepam group (upper end of the 95% confidence interval at 6.28 bpm above baseline). During the second hour after drug administration, this decrease in heart rate diminished overall, and the change compared to baseline values was even mildly positive (change from 0.23 to 5.85 bpm) in animals of all three test groups (6/10 midazolam, 4/10 in 2-mg/kg tiletamine-zolazepam, and 2/10 in 3- mg/kg tiletamine-zolazepam group). Once righting reflexes were regained (21.0–99.4 min depending on anesthetic group, Table 2), heart and respiratory rates had returned to baseline values in all snakes.

Measured respiratory rate patterns were similar to heart rate patterns. During the first hour after drug administration, animals of all three drug regimen groups experienced slight decreases in breaths taken per minute (breaths/min), whereas some animals experienced small increases (4/10 animals of the midazolam group, 4/10 of the 2-mg/kg tiletamine-zolazepam group, and 6/10 of the 3-mg/kg tiletamine-zolazepam group). In midazolam-treated snakes, the mean change from baseline values ranged from -0.85 to -6.65 breaths/min and diminished during the second hour after drug administration to +0.85 to -2.65 breaths/min (Table 3). Snakes exposed to 2 mg/kg of tiletamine+zolazepam experienced a mean change of +1.5 to -5.7 breaths/min during the first hour of drug administration and +1.2 to -3.8 breaths/min during the second hour. These changes in respiratory rates were not significantly different from those of animals in the midazolam-treated group. However, snakes injected with 3 mg/kg of tiletamine+zolazepam experienced a mean change of +2.4 to -2.3 breaths/min during the first hour of drug administration. This change was statistically significantly different from the minimum mean respiratory rate change at baseline when compared to the 2-mg/kg of tiletamine+zolazepam and midazolam groups (Table 3). During the second hour of drug administration, respiratory rates of these same animals dropped to a mean change of -0.5 to -6.0 bpm compared to baseline values.

## DISCUSSION

Anesthetic regimens for reptiles can be grouped into inhaled compounds, injectable compounds, or a combination of both. Administration of inhaled compounds is challenging since reptiles can hold their breaths for various durations, and intubation is therefore required to ensure control of drug administration and prevent hypoxia [53-56]. Intubation adds an additional step that requires veterinary expertise and additional equipment that needs to be added to the laboratory setting. This step complicates study approval involving infectious agent, i.e. when the safety of the laboratory worker is a primary concern. Therefore, injectable compounds would be advantageous for reptile research in containment settings. However, these compounds can vary widely in their effectiveness and may be associated with prolonged recoveries. For instance, the widely used *N-*methyl-D-aspartic acid (NMDA) antagonist ketamine is associated with prolonged recoveries, poor analgesia, and apnea in snakes when used alone [45, 57]. The addition of alpha2 adrenergic agonists (e.g., dexmedetomidine) to ketamine-based anesthesia improves the quality of the anesthesia and reduces recovery time [45], but in the authors’ experience still results in cardiovascular depression that may warrant assisted ventilation and supplemental oxygen. Propofol is associated with fast, smooth induction of anesthesia but must be administered via intravenous or intracardiac routes and is therefore a challenge to use in snakes [55, 56]. Alfaxalone or alfaxalone/alphadalone combinations are a challenge to use effectively in some reptiles. Recent work using ball pythons demonstrated smooth and rapid induction of anesthesia after intramuscular administration—however, apnea was still observed at higher doses [58-62].

Midazolam, a benzodiazepine sedative, has proven effective for restraint of some reptiles, but generally is not well studied [45-52]. In snakes, midazolam is commonly used in conjunction with ketamine for induction of anesthesia [45]. Tiletamine+zolazepam is widely used for mammal anesthesia, immobilization, and biosample collection, however peer reviewed and published literature providing detailed descriptions of the effects of tiletamine+zolazepam on reptiles and amphibians is scant [40-43]. However, high doses of tiletamine+zolazepam over 20 mg/kg achieved anesthesia in boa constrictors [42, 44]. The administration of a fixed combination of tiletamine (dissociative anesthetic) and zolazepam (benzodiazepine sedative), at doses above 6 mg/kg is associated with prolonged recoveries in reptiles. However at lower doses, 2–4 mg/kg, the combination is useful for restraint and induction of short-term anesthesia [45]. The main goal of this study was to identify an injectable compound or compound combination that could be used to achieve anesthesia in ball pythons for future research in an infectious disease setting. We compared the quality and duration of sedation and anesthesia produced by intramuscular midazolam (2 mg/kg) and tiletamine+zolazepam (2 mg/kg and 3 mg/kg) in healthy ball pythons. No clinical abnormalities (e.g., pale mucous membranes, irregular heartbeat, apnea) were noted in any animal after drug administration. Snakes of all three drug groups had temporary mild-to-moderate depression of heart rate and respiratory rate not severe enough to require intubation. This depression has also been reported for various mammals exposed to tiletamine+zolazepam [63-65]. On the other hand, an increase in heart rate, approximately 10 bpm above baseline, was measured in boa constrictors and in dogs [42, 44, 66]. In our study, an increase in heart rate of this magnitude was observed in only two ball pythons, both of which received 2 mg/kg tiletamine+zolazepam. In boa constrictors, tiletamine+zolazepam also results in respiratory rate decreases similar to those observed in our study [42, 44]. However, the dose of tiletamine+zolazepam combination used in the boa constrictor study (25 mg/kg of metabolic body weight administered intramuscularly; adjusted 21 mg/kg based on allometric scaling and metabolic body weight) was considerably higher than those used in our study (2 and 3 mg/kg) [45]. These findings indicate tiletamine+zolazepam has species-specific effects.

The effectiveness of midazolam as a sedative for reptiles varies widely depending on which reptile is exposed. The accepted dose for snakes is 1–2 mg/kg intravenously or intramuscularly [45]. Since midazolam is a sedative rather than a true anesthetic, we predicted that this drug would not result in anesthesia in ball pythons. In our experiments, midazolam administration was associated with the same induction time as the tiletamine+zolazepam combinations. However, as expected, the efficacy of midazolam in immobilizing ball pythons was poor (n = 6/10). Midazolam was also associated with the fastest recovery time of all tested drug regimens. We expected that not all midazolam-treated snakes would lose their righting reflexes and enter Stage 2 of anesthesia, and indeed the number of snakes failing to achieve this stage of anesthesia was notable (n = 4/10). An unexpected finding was that one midazolam-treated animal failed to achieve any level of sedation (Table 1, Figure 1). Importantly, the injection volume required for administration of 2 mg/kg of midazolam was, on average, 10 times the volumes used for administering the tiletamine+zolazepam combinations (for a 1.5-kg animal, dose volumes were 0.6 ml vs. 0.03 ml or 0.04 ml of tiletamine+zolazepam). For ball pythons in our study weighing over 1.2 kg, the midazolam dose volume surpassed 0.5 ml, which is considered too large of a volume to accurately administer intramuscularly [67].

The commonly accepted dosage range for tiletamine+zolazepam in snakes is 2–6 mg/kg, although protracted recovery times of 48 to 72 hours have been observed by clinical veterinarians using the upper end of this dose range [45]. We observed clear differences in the quality of anesthesia between the 2-mg/kg and 3-mg/kg tiletamine+zolazepam groups. In our study, using the 2-mg/kg dose resulted in reliable and smooth sedation and good snake immobilization of sufficient duration for further manipulation (57.7 ± 30.5 min) in 7 of 10 animals. Recovery periods were less than 1.5 hours. However, the 2-mg/kg dose did not result in Stage 3 of anesthesia in 7 of 10 animals. On the other hand, the 3-mg/kg dose yielded more promising results, with all animals entering Stage 2 and 3 of anesthesia. Loss of the withdrawal reflex (Stage 3 of anesthesia) lasted only 10–30 min. This time frame suffices only for minor manipulations such as pathogen exposure or blood sample collection.

A drawback to the 3-mg/kg dose of tiletamine+zolazepam is the significant increase in length of snake recovery times compared to those observed for snakes treated with 2 mg/kg of tiletamine+zolazepam or 2 mg/kg of midazolam. Recovery times were widely variable among different snakes administered the drug combination at 3mg/kg as demonstrated by the large 95% confidence limit and standard deviation for both the time to regain righting reflex (mean 94.4 min) and time to resume normal behavior (median of 3.75 hours and mean of 5.85 hours. (Table 2). This highly variable recovery time and lingering sedation (as well as the variable induction length) between individual snakes receiving 3 mg/kg of tiletamine+zolazepam may be due to differing pharmacokinetic clearance mechanisms of tiletamine and zolazepam. In pigs, tiletamine is metabolized much faster than zolazepam, and zolazepam metabolites retain more pharmacologic activity than tiletamine metabolites, thus prolonging the sedative effect of the drug combination beyond its dissociative effects [68].

Pharmacokinetic data are not available for tiletamine or zolazepam in reptiles, presenting a potential topic for future research. It also remains to be determined whether reptiles metabolize tiletamine and zolazepam primarily through the kidneys as has been demonstrated for mammals [68]. This lack of pharmacokinetic data and pKa studies in reptiles presents a considerable limitation to our study and other investigations of the performance of anesthetic compounds in reptiles. Without definite knowledge of the half-life of midazolam, tiletamine, and zolazepam in ball pythons, we can only surmise that when the snakes resumed normal behavior, they had most likely cleared sufficient amounts of the drugs to reduce blood concentration below therapeutic concentrations.

In conclusion, 3 mg/kg of intramuscular tiletamine+zolazepam is an adequate drug regimen to achieve short-term immobilization in ball pythons. Future studies should confirm these findings with a higher number of animals, a wider range of doses, and more invasive anesthetic monitoring, such as blood pressure, to facilitate more direct comparisons to experimental settings with similar work already performed with boa constrictors [42, 44].

## ACKNOWLEDGEMENTS

We thank Laura Bollinger (IRF-Frederick) for critically editing this manuscript and Jiro Wada for medical illustration and graphic support (IRF-Frederick). We thank the SEAVS technical staff (Jennifer Hutchins, LVT) for her assistance in animal procurement and quarantine. We thank Scott Stahl, DVM, Emily Nielsen, DVM, and Outback Reptiles (Manassas, VA, USA) for donation of the animals. The findings and conclusions in this report are those of the authors and do not necessarily reflect the views or policies of the US Department of the Army, the US Department of Defense, the US Department of Health and Human Services or of the institutions and companies affiliated with the authors.

